# Minimal phrase composition revealed by intracranial recordings

**DOI:** 10.1101/2021.04.30.442171

**Authors:** Elliot Murphy, Oscar Woolnough, Patrick S. Rollo, Zachary Roccaforte, Katrien Segaert, Peter Hagoort, Nitin Tandon

## Abstract

The ability to comprehend phrases is an essential integrative property of the brain. Here we evaluate the neural processes that enable the transition from single word processing to a minimal compositional scheme. Previous research has reported conflicting timing effects of composition, and disagreement persists with respect to inferior frontal and posterior temporal contributions. To address these issues, 19 patients (10 male, 19 female) implanted with penetrating depth or surface subdural intracranial electrodes heard auditory recordings of adjective-noun, pseudoword-noun and adjective-pseudoword phrases and judged whether the phrase matched a picture. Stimulus-dependent alterations in broadband gamma activity, low frequency power and phase-locking values across the language-dominant left hemisphere were derived. This revealed a mosaic located in the posterior superior temporal sulcus (pSTS), in which closely neighboring cortical sites displayed exclusive sensitivity to either lexicality or phrase structure, but not both. Distinct timings were found for effects of phrase composition (210–300 ms) and pseudoword processing (approximately 300–700 ms), and these were localized to neighboring electrodes in pSTS. The pars triangularis and temporal pole encode anticipation of composition in broadband low frequencies, and both regions exhibit greater functional connectivity with pSTS during phrase composition. Our results suggest that the pSTS is a highly specialized region comprised of sparsely interwoven heterogeneous constituents that encodes both lower and higher level linguistic features. This hub in pSTS for minimal phrase processing may form the neural basis for the human-specific computational capacity for forming hierarchically organized linguistic structures.

**Significance:** Linguists have claimed that the integration of multiple words into a phrase demands a computational procedure distinct from single word processing. Here, we provide intracranial recordings from a large patient cohort, with high spatiotemporal resolution, to track the cortical dynamics of phrase composition. Epileptic patients volunteered to participate in a task in which they listened to phrases (“red boat”), word-pseudoword or pseudoword-word pairs (e.g., “red fulg”). At the onset of the second word in phrases, greater broadband high gamma activity was found in posterior superior temporal sulcus in electrodes that exclusively indexed phrasal meaning, and not lexical meaning. These results provide direct, high-resolution signatures of minimal phrase composition in humans, a potentially species-specific computational capacity.

## Introduction

How the brain integrates individual word meanings to comprehend multi-word utterances is an issue that has vexed the cognitive neuroscience of language for decades (Hagoort, 2020). This linguistic compositional process – the combination of words into larger structures with new and complex meaning – has been referred to as “Merge” (Chomsky et al., 2019) or “Unification” (Hagoort, 2013). A simple paradigm for studying complex meaning is the use of minimal phrases, such as in the *red-boat* paradigm, which focuses on simple combinations of two words and avoids confounds associated with more complex linguistic stimuli (Bemis and Pylkkänen, 2011, 2013; Brennan and Pylkkänen, 2012; Bozic et al., 2015; Flick et al., 2018; Flick and Pylkkänen, 2020; Pylkkänen, 2020). A “red boat” is interpreted as a boat which is red, and not a red object which hosts boat-related properties, with phrases delivering novel syntactic and conceptual formats (Murphy, 2015). *Red-boat* experiments isolate semantic composition, which in turn encompasses syntactic composition.

Functional neuroimaging studies using the *red-boat* paradigm implicate the left anterior temporal lobe (ATL), specifically the temporal pole (Antonucci et al., 2008; Lambon Ralph et al., 2012; Wilson et al., 2014; Zhang and Pylkkänen, 2018), inferior frontal regions (Graessner et al., 2021b, 2021a) and posterior temporal regions (Flick and Pylkkänen, 2020; Matchin and Hickok, 2020) as crucial nodes for phrase composition with variations in the timing (180– 350ms post-composition) and duration (50–100ms) of their engagement (Kochari et al., 2021). Posterior temporal cortex is more broadly implicated in syntactic comprehension and production (Duffau et al., 2014; Artoni et al., 2020; Matchin and Hickok, 2020; Graessner et al., 2021b; Lopopolo et al., 2021). The left posterior temporal lobe and angular gyrus (AG) show greater activity for sentences than word lists, and for phrases than words, making them candidate regions for the retrieval of phrasal templates (Hagoort, 2003, 2017; Brennan et al., 2016; Matchin and Hickok, 2020). According to a range of parsing models, adjective-noun syntax is also constructed *predictively* (Berwick et al., 2019), and anticipatory amplitude increases have been isolated to the alpha/beta bands (Gastaldon et al., 2020).

Phrase composition is a rapid process that is likely dependent on finely organized sets of distributed cortical substrates. Previous work using the *red-boat* paradigm has been limited by spatiotemporal resolution. Using recordings from intracranial electroencephalography (iEEG) with depth electrodes penetrating grey matter or electrodes located on the cortical surface, we conducted a study of minimal phrase composition with auditory presentations of the red boat paradigm, building directly and using stimuli from an established design (Bemis and Pylkkänen, 2013). Given our large cohort, we were able to perform a data driven analysis of iEEG data from the whole brain acquired at unprecedented spatiotemporal resolution. Most evidence that left ATL and posterior temporal regions are involved in basic composition is from magnetoencephalography (MEG), while the evidence supporting a role for IFG mostly comes from functional magnetic resonance imaging (fMRI). Signal loss in fMRI may partly explain the lack of ATL effects in this portion of the imaging literature (due to proximity to sinuses) (Olman et al., 2009; Bonner and Price, 2013), yet even within the MEG literature there is variation in the timing of composition effects. This suggests that intracranial recordings can contribute to resolving the spatiotemporal dynamics of phrase processing. Our central research questions concerned: (i) the precise localization of composition effects, addressing a current tension in the literature with respect to frontal, temporal and parietal contributions to minimal phrase processing (Schell et al., 2017; Flick and Pylkkänen, 2020; Graessner et al., 2021a); (ii) the precise timing and duration of composition effects, addressing the above noted variations documented in the literature.

## Materials and Methods

### Participants

19 patients (10 male, 18-41 years, IQ 97 ± 12, 2 left-handed) participated in the experiment after written informed consent was obtained. All were native English speakers. All experimental procedures were reviewed and approved by the Committee for the Protection of Human Subjects (CPHS) of the University of Texas Health Science Center at Houston as Protocol Number HSC-MS-06-0385.

### Electrode Implantation and Data Recording

Data were acquired from either subdural grid electrodes (SDEs; 6 patients) or stereotactically placed depth electrodes (sEEGs; 13 patients) (Fig. 1B). SDEs were subdural platinum-iridium electrodes embedded in a silicone elastomer sheet (PMT Corporation; top-hat design; 3mm diameter cortical contact), and were surgically implanted via a craniotomy (Conner et al., 2011; Pieters et al., 2013; Tong et al., 2020; Kohlhase et al., 2021). sEEGs were implanted using a Robotic Surgical Assistant (ROSA; Medtech, Montpellier, France) (Rollo et al., 2020; McCarty et al., 2021). Each sEEG probe (PMT corporation, Chanhassen, Minnesota) was 0.8mm in diameter and had 8-16 electrode contacts. Each contact was a platinum-iridium cylinder, 2.0mm in length and separated from the adjacent contact by 1.5-2.43mm. SDE patients had a number of cortical arrays implanted (mean ± SD: 9 ± 1) with a mean of 146.5 electrodes (SD ± 53.5). sEEG patients had penetrating depth probes (mean 15.1 ± 2.3) with a mean of 199.9 electrodes (SD ± 30.1). Typical coverage was fronto-temporal, dictated by location of the epilepsy in the antero-mesial temporal lobe in the majority, with parietal and occipital coverage in a number of patients. Following surgical implantation, electrodes were localized by co-registration of pre-operative anatomical 3T MRI and post-operative CT scans in AFNI (Cox, 1996). Electrode positions were projected onto a cortical surface model generated in FreeSurfer (Dale et al., 1999), and displayed on the cortical surface model for visualization (Pieters et al., 2013). Intracranial data were collected during research experiments starting on the first day after electrode implantation for sEEGs and two days after implantation for SDEs. Data were digitized at 2 kHz using the NeuroPort recording system (Blackrock Microsystems, Salt Lake City, Utah), imported into Matlab, initially referenced to the white matter channel used as a reference for the clinical acquisition system and visually inspected for line noise, artifacts and epileptic activity. iEEG provides uniquely high spatiotemporal resolution recordings and is less susceptible to artifacts (e.g., muscle movements) (Flinker et al., 2011; Arya, 2019). Electrodes with excessive line noise or localized to sites of seizure onset were excluded. Each electrode was re-referenced to the common average of the remaining channels. Trials contaminated by inter-ictal epileptic spikes, saccade artefacts and trials in which participants responded incorrectly were discarded. Electrodes contributing to regions of interest were taken from both SDE and sEEG patients; for instance, we follow previous intracranial work that has included subdural contacts in monitoring activity from posterior temporal sulcus (Uno et al., 2015).

**Figure 1:**
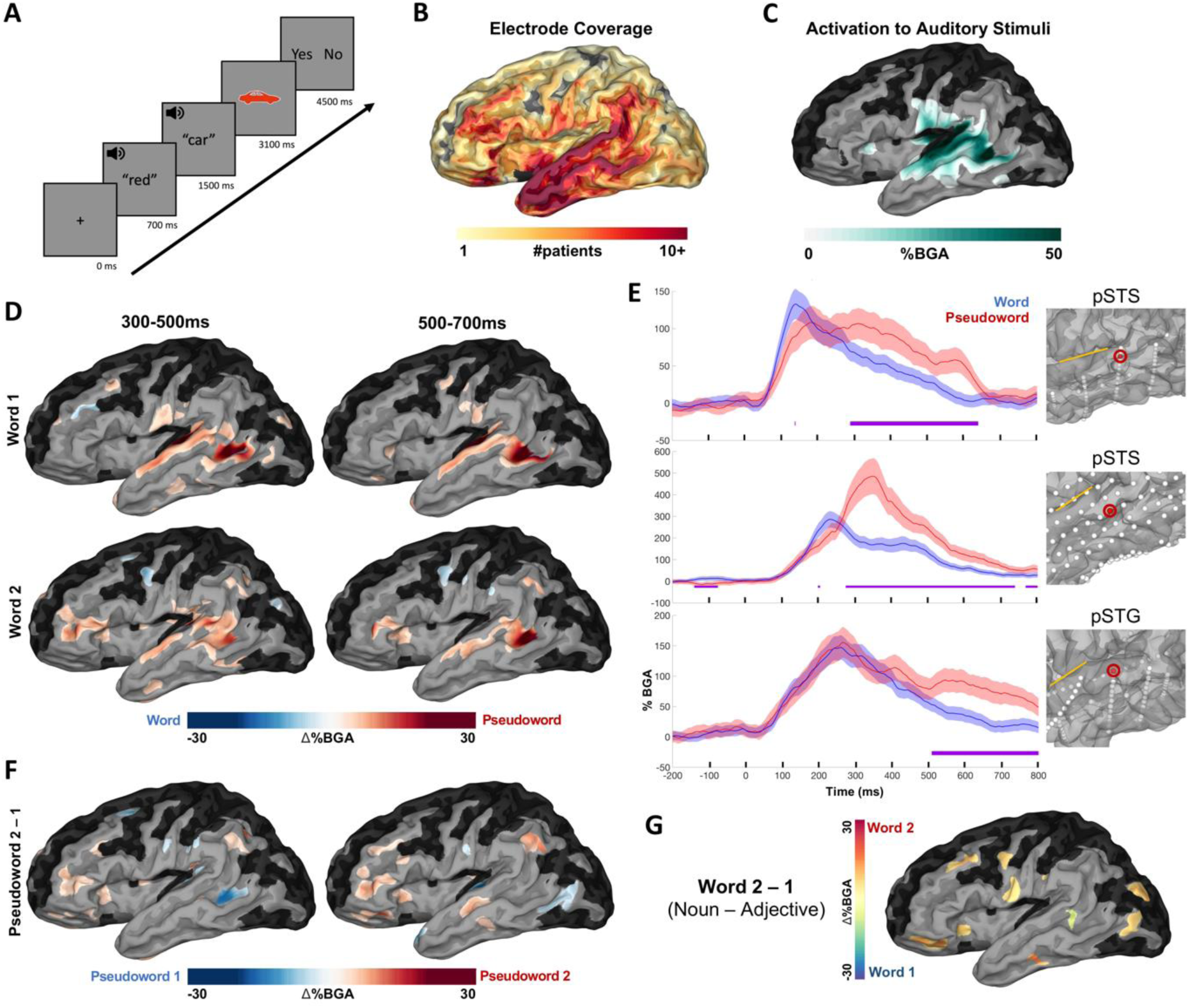
Patient coverage map and grouped analysis for lexicality. (A) Experimental design. Average stimuli length: adjectives (420 ± 39 ms; mean ± SD), nouns (450 ± 75 ms), pseudowords (430 ± 38 ms). (B) Group coverage map of left hemisphere electrodes included in analyses, plotted on a semi-inflated standardised N27 surface. (C) Broadband gamma activity (BGA) increases from pre-stimulus baseline (−500 to -100ms prior to first word) for all conditions from 100 to 400 ms after first word onset (threshold: %BGA > 5%, t > 1.96, patient coverage ≥ 3; *p* < 0.01 corrected). Black surfaces fell below patient coverage threshold. (D) SB-MEMA comparing words vs pseudowords. Red coloration indexes greater BGA (70–150 Hz) for pseudowords, and blue for words (same thresholds as in (C)). Top row: word position 1; bottom row: word position 2. (E) Exemplar electrodes for the words vs pseudowords analysis. Error bars (colored shading) set at one standard error. Sylvian fissure is marked with a yellow line for reference on each surface. Time 0 ms = word 1 onset. (F) SB-MEMA indicating BGA increases for pseudowords at the second word position relative to pseudowords at the first word position (time windows collapsed for both word 1 and word 2 positions). (G) SB-MEMA contrast for real words from both non-compositional conditions across the 300–500 ms window (Adjective-Pseudoword and Pseudoword-Noun). Colors indicate nouns (red) and adjectives (blue).

### Experimental Design and Statistical Analysis

Grammatical noun phrases (“red boat”) and pseudoword phrases with variable positions of the pseudoword (“bleeg boat”, “red fulg”) were used to isolate semantic compositional processing. An adjective-pseudoword condition (“red fulg”) allowed for the isolation of semantic compositionality and reduced predictability. The inclusion of pseudowords in our stimulus set builds upon a previous study, from which we abridged our list of words and graphic images (Bemis and Pylkkänen, 2013). We focused on high frequency gamma changes (Forseth et al., 2018; Conner et al., 2019; Johnson et al., 2020; Leszczyński et al., 2020) that typically index local cortical processing and are implicated in a range of cognitive processes (Buzsáki and Watson, 2012; Hovsepyan et al., 2020; Packard et al., 2020). An early anterior negative deflection immediately preceding the critical noun in combinatorial contexts (−50–100ms) likely indexes syntactic prediction (Neufeld et al., 2016). Low frequency power increases have also been noted during the anticipatory window and during phrase composition (Bastiaansen and Hagoort, 2015; Lewis et al., 2016; Segaert et al., 2018), and in a variety of auditory phrase and sentence processing paradigms (Ding et al., 2016; Mai et al., 2016; Keitel et al., 2017, 2018; Leivada and Murphy, 2021). Therefore, we also evaluated the role of low frequencies during anticipatory composition, focusing on the alpha/beta bands given the joint involvement of these in the literature and the similar anticipatory dynamics attributed to them. Lastly, we asked patients to determine whether the words they heard matched a subsequent image, enabling validation of their attention as well as analyses related to phrase-matched and phrase-contrasted contexts. Our analyses were restricted to language-dominant left hemisphere electrode coverage.

Participants were presented with two-word auditory phrases, grouped by three conditions: Adjective-Noun (“red boat”), Adjective-Pseudoword (“red neub”), Pseudoword-Noun (“zuik boat”). Since these pseudoword phrases include phonologically viable nonwords, differences in the second position of the phrase between these items and the grammatical noun phrases are likely a result of compositional processing. While most previous studies have presented a licensable noun, our inclusion of the Adjective-Pseudoword condition further isolates composition and reduces predictability. To ensure attention was maintained, after each trial participants were shown a colored drawing and asked to press a button indicating whether the picture matched the phrase they had just heard. Participants were told to respond positively only when the picture fully matched the phrase. Auditory and visual stimuli were adapted from a previous *red-boat* experiment (Bemis and Pylkkänen, 2013), from which we obtained our list of real words and graphic images.

A fixation cross was presented in the centre of the screen for 700 ms followed by the first word, and 800 ms later the second word was presented. 1600 ms after the onset of the second word the picture was presented, and 1400 ms after picture presentation participants were prompted to respond (Fig. 1A). Following their response, a blank screen was shown for 1500 ms. Stimuli were presented in a pseudorandom order, with no repetition amongst items. The number of trials per block across the full experiment was as follows: Adjective-Noun (80), Pseudoword-Noun (40), Adjective-Pseudoword (40). All patients undertook 2 blocks. Half of the Adjective-Noun trials matched the picture presented (i.e., “red boat” was heard by the patient, and a picture of a red boat was then presented), and the other half did not match. 6 adjectives were used: black, blue, brown, green, pink, red (length M: 4.3, SD: 0.7; SUBTLEXus log-frequency 3.64). 20 nouns were used: bag, bell, boat, bone, cane, cross, cup, disc, flag, fork, hand, heart, house, key, lamp, leaf, lock, plane, shoe, star (length M: 4.0, SD: 0.6; log-frequency 3.38) (Brysbaert et al., 2012). 6 pseudowords were used: beeg, cresp, kleg, nar, neub, zuik (length M: 4.0, SD: 0.6). Average stimuli length: Adjectives (420 ms), Nouns (450 ms), Pseudowords (430ms). Stimuli were presented using Psychtoolbox (Kleiner et al., 2007) on a 15.4” LCD screen positioned at eye-level, 2-3’ from the patient. Auditory stimuli were presented using stereo speakers (44.1 kHz, MacBook Pro 2015).

A total of 3458 electrode contacts were implanted in patients; 2135 of these were included for analysis after excluding channels proximal to the seizure onset zone or exhibiting excessive inter-ictal spikes or line noise. Analyses were performed by first bandpass filtering the raw data of each electrode into broadband gamma activity (BGA; 70–150 Hz) following removal of line noise and its harmonics (zero-phase 2nd order Butterworth band-stop filters). Electrodes were also visually inspected for saccade artefacts. A frequency domain bandpass Hilbert transform (paired sigmoid flanks with half-width 1.5 Hz) was applied and the analytic amplitude was smoothed (Savitzky-Golay FIR, 3rd order, frame length of 251ms; Matlab 2019b, Mathworks, Natick, MA). BGA was defined as percentage change from baseline level; -500 to -100ms before the presentation of the first word in each trial. Periods of significant activation were tested using a one-tailed t-test at each time point and were corrected for multiple comparisons with a Benjamini-Hochberg false detection rate (FDR) threshold of q<0.05, where the q-value denotes the standard term given to adjusted p-values for FDR significance corrected for multiple comparisons (Benjamini and Hochberg, 1995; Storey, 2011). For the grouped analysis, all electrodes were averaged within each subject and then the between-subject averages were used.

To provide statistically robust and topologically precise estimates of BGA, and to account for variations in sampling density, population-level representations were created using surface-based mixed-effects multilevel analysis (SB-MEMA) (Fischl et al., 1999; Conner et al., 2011; Kadipasaoglu et al., 2014, 2015). This method accounts for sparse sampling, outlier inferences, as well as intra- and inter-subject variability to produce population maps of cortical activity. A geodesic Gaussian smoothing filter (3mm full-width at half-maximum) was applied. Significance levels were computed at a corrected alpha-level of 0.01 using family-wise error rate corrections for multiple comparisons. The minimum criterion for the family-wise error rate was determined by white-noise clustering analysis (Monte Carlo simulations, 5000 iterations) of data with the same dimension and smoothness as that analyzed (Kadipasaoglu et al., 2014). Results were further restricted to regions with at least three patients contributing to coverage and BGA percent change exceeding 5%.

Anatomical groups of electrodes were delineated, firstly, through indexing electrodes to the closest node on the standardized cortical surface (Saad and Reynolds, 2012), and secondly, through grouping channels into parcellations determined by Human Connectome Project (HCP) space (Glasser et al., 2016). Parametric statistics were used since HCP regions of interest contained over 30 electrodes. When contrasting experimental conditions, two-sided paired t-tests were evaluated at each time point for each region and significance levels were computed at q < 0.01 using an FDR correction for multiple comparisons.

To generate event-related potentials (ERPs), the raw data were band pass filtered (0.1–50 Hz). Trials were averaged together and the resultant waveform was smoothed (Savitzky-Golay FIR, third-order, frame length of 251 ms). To account for differences in polarity between electrodes, ERPs were converted to root mean square (RMS), using a 50 ms sliding window. All electrodes were averaged within each subject, within ROI, and then the between-subject averages were used.

To explore the functional connectivity between ROIs, we used a generalized phase-locking analysis to estimate the dominant spatiotemporal distributions of field activity, and the strength of the coupling between them. Phase information was extracted from the down-sampled (200 Hz) and wide band-pass filtered data (3–50 Hz; zero-phase 8th order Butterworth band-pass filter) using the ‘generalized phase’ method (Davis et al., 2020) with a single-sided Fourier transform approach. This method captures the phase of the predominant fluctuations in the wideband signal and minimizes filter-related distortion of the waveform. PLV was calculated as the circular mean (absolute vector length) of the instantaneous phase difference between each electrode pair at each time point and baselined to the pre-stimulus period -500 to -100 ms before onset of the first word. Statistics were calculated using the mean PLV of correctly answered trials between 0 to 500 ms after second word onset, comparing against a null distribution generated by randomly re-pairing trial recordings across the electrode pairs 500 times. Significant PLV from pre-stimulus baseline was accepted at a threshold of p < 0.05. When computing conditional differences, significance was accepted at q < 0.05 using an FDR correction for multiple comparisons.

### Data Accessibility

The datasets generated from this research are not publicly available due to their containing information non-compliant with HIPAA, and the human participants from whom the data were collected have not consented to their public release. However, they are available on request from the corresponding author.

### Code Accessibility

The custom code that supports the findings of this study is available from the corresponding author on request.

## Results

Patients were presented with auditory recordings grouped randomly into three conditions: Adjective-Noun, Pseudoword-Noun, Adjective-Pseudoword. A subsequent colored image was presented, and patients were tasked with responding, with a button press, to whether the image matched the phrase or not (Fig. 1A). Across the cohort, we had good coverage over lateral and medial temporal lobe, inferior parietal lobe and inferior frontal regions, with some coverage reaching into other portions of fronto-parietal cortex (Fig. 1B). We saw activation in response to the auditory stimuli most prominently in superior and middle temporal regions (Fig. 1C). Below we report results pertaining to lexicality; phrase anticipation; phrase composition; and linguistic-visual unification.

### Behavioral performance

Performance in the image matching task was highly accurate at 97 ± 3% (311 ± 11/320 trials), with an average response time of 1599 ± 539 ms. Only correct trials were analyzed further.

### Effects of lexicality

To disentangle single word semantic effects from those of combinatorial semantics, we probed the difference in representation between words and pseudowords. We generated a surface-based, population-level map of cortical activity using a surface-based mixed-effects multi-level analysis (SB-MEMA) (Fischl et al., 1999; Conner et al., 2011; Kadipasaoglu et al., 2014, 2015), a method specifically designed to account for sampling variations in iEEG and minimize effects of outliers. An SB-MEMA contrasting adjectives and pseudowords in word position 1, and nouns and pseudowords in word position 2 (Fig. 1D), revealed significantly greater broadband gamma activity (BGA; 70–150 Hz) for pseudowords than words in posterior superior temporal gyrus (pSTG) (word 1, 300–500 ms: *β* = 0.88; *p* = .006), posterior superior temporal sulcus (pSTS) (word 1, 300–700 ms: *β* = 0.30; *p* <.001; word 2, 300–700 ms: *β* = 0.23; *p* <.001) and pars triangularis (word 2, 300–700 ms: *β* = 0.08; *p* = .004). We found greater BGA increases for pseudowords than words at position 2 than position 1 around pars triangularis and surrounding frontal areas (Fig. 1F) (pars triangularis, 300–700 ms: *β* = 0.09; *p* <.001; pSTS, 300–500 ms: *β* = 0.09; *p* = .008). A Word 2–1 subtraction for both non-compositional trials (Adjective-Pseudoword, Pseudoword-Adjective) revealed no such effects in posterior temporal regions, and a BGA increase for nouns over frontal opercular sites (300–500 ms: *β* = 0.10; *p* = .002) (Fig. 1G).

### Compositional anticipation

We next contrasted Adjective-Noun and Pseudoword-Noun conditions at the onset of second word presentation, with only the former condition licensing any phrasal anticipation. Given that both traditional alpha (8–12 Hz) and beta (12–30 Hz) bands have been regularly implicated in linguistic prediction and anticipatory composition (Lewis et al., 2016; Segaert et al., 2018; Hardy et al., 2021), with previous research in this domain (Gisladottir et al., 2018; Terporten et al., 2019) and neighboring domains (Piai et al., 2020) collapsing these bands, we analyzed activity across the 8–30 Hz range. During the anticipatory window for phrase formation (from -200ms to 0 ms prior to the second word onset), low frequency power (8–30 Hz) exhibited a significant conditional difference (Fig. 2). Greater alpha/beta power for anticipatory trials was found in pars triangularis (*β* = 0.18; *p* = .009), ATL (*β* = 0.15; *p* = .008) and STG (*β* = 0.13; *p* = .008). These effects were unrelated to BGA. For comparison, we also plot the anticipatory window in BGA for -100 to 0 ms (Fig. 2B), which exhibited no clear relation with the lower frequency effects.

**Figure 2:**
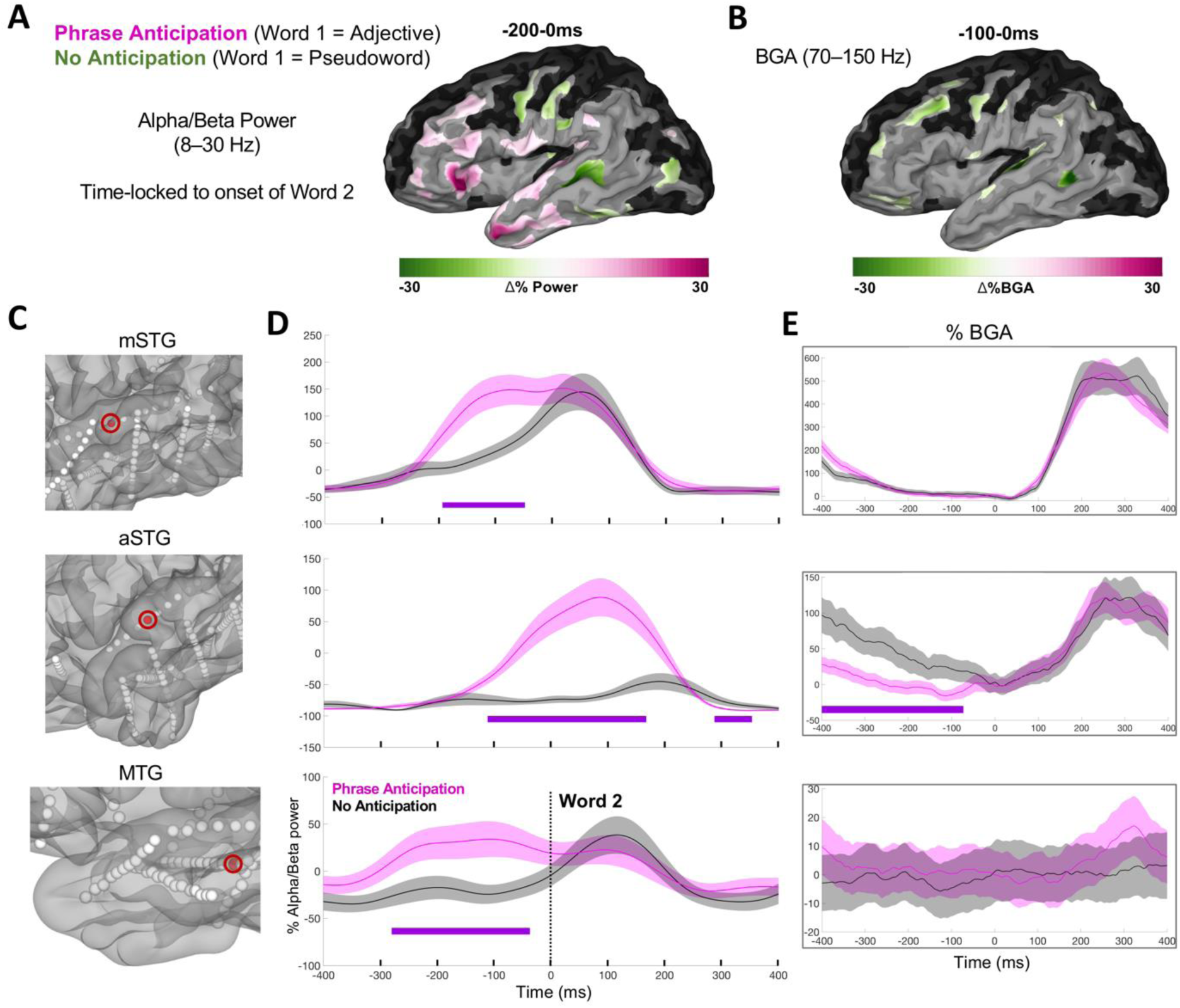
Syntactic-semantic compositional anticipation. (A) SB-MEMA for the anticipatory time window centred around second word onset (−200–0 ms) for low frequency alpha/beta power. Pink: Greater power for composition anticipation. Green: Greater for no anticipation (i.e., after having heard a Pseudoword at word 1 position). The same SB-MEMA thresholds as in Fig. 1C were applied. (B) SB-MEMA for BGA for -100–0 ms prior to word 2 onset. (C)-(E) Exemplar electrodes with location (C), low frequency power traces (D) and BGA traces (E) across three patients sorted by row.

### Phrase composition

The combinatorial contrast (Adjective-Noun vs [Adjective-Pseudoword + Pseudoword-Noun]) revealed greater BGA for portions of pSTS during phrase composition than non-composition (100–300 ms: *β* = 0.10; *p* = .003) (Fig. 3A). The specific onset of this effect was around 210ms after noun onset, with peak BGA at around 300ms (Fig. 4B). The same region exhibited greater BGA for non-compositional trials at later time points (300–700 ms: *β* = 0.10; *p* = .001). Certain portions of pSTS across patients displayed exclusive sensitivity to phrase composition, and not lexicality (Fig. 3B, 3C). Other regions – pSTG and inferior frontal gyrus (IFG) – did not show any significant BGA differences between these conditions (Fig. 4A). ERP responses were dissociable from BGA across the ROIs plotted: A late effect for non-composition was found in Broca’s area, and a late signature was detected in temporal pole for composition.

**Figure 3:**
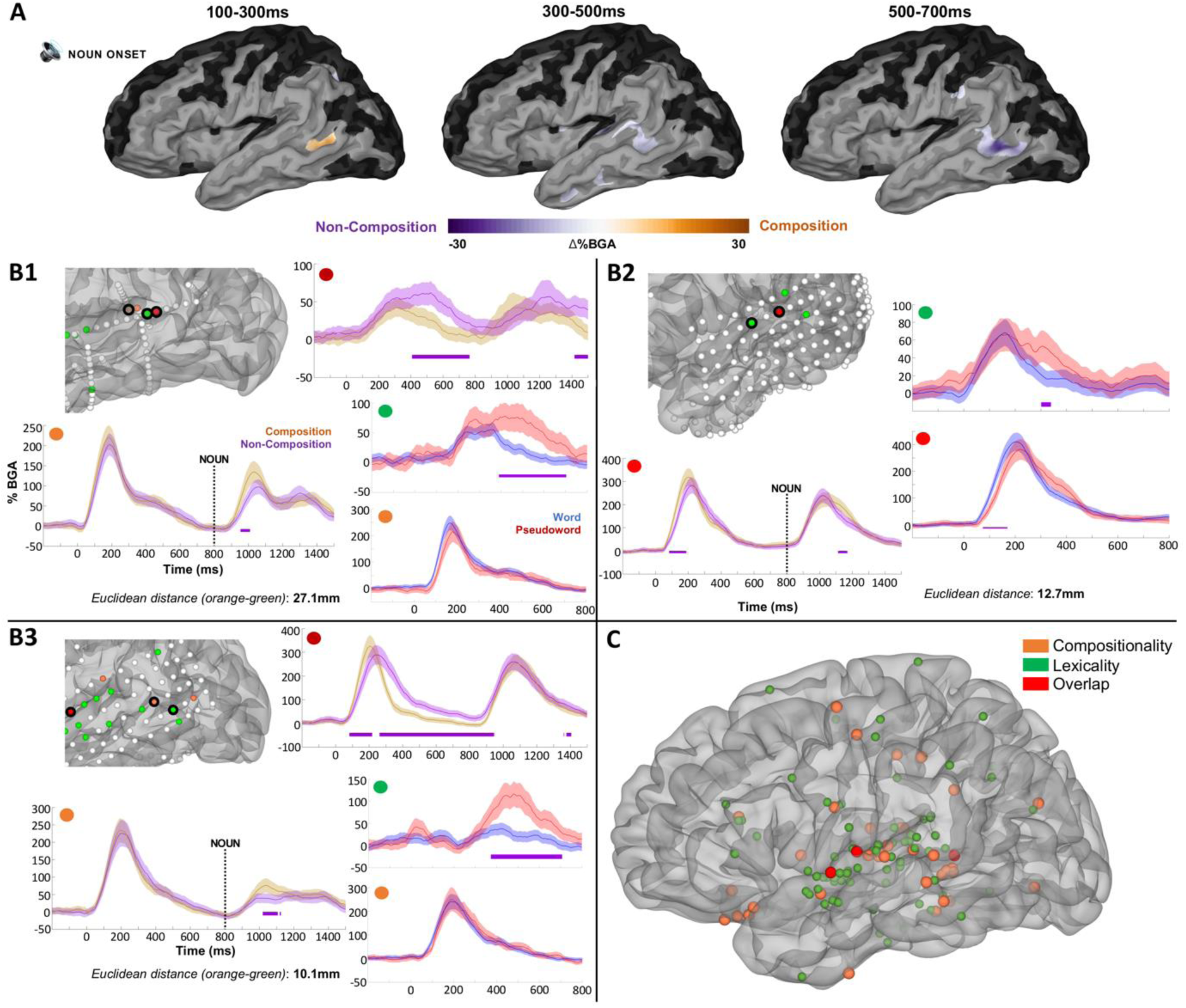
Grouped analysis for phrase composition. (A) SB-MEMAs for the phrase composition analysis for BGA (70–150 Hz). Orange indexes greater BGA for phrase composition and purple indexes greater BGA for non-composition. The same SB-MEMA thresholds as in Fig. 1C were applied. Time 0 ms = word 2 onset. (B) Exemplar electrodes with FDR-corrected (one-tailed t-tests, q < 0.05) significance bars in purple plotted in native patient space. Includes compositional contrast and lexicality contrast, dissociating neighboring portions of pSTS that responded exclusively for phrase composition and not lexicality (orange dots = composition effect; green dots = lexicality effect; red dots = effect for both composition and lexicality). Electrodes that showed greater, later BGA increases for non-composition were spatially distinct from those sensitive to composition in early windows. Error bars set at one standard error. Numbers denote distinct patients (B1–B3). Time 0ms = word 1 onset; word 2 onset at 800ms. (C) Electrodes exhibiting a significant BGA contrast for compositional vs non-compositional trials, and words vs pseudowords. 39 electrodes (orange) from 12 patients exhibited an FDR-significant contrast (one-tailed t-tests, q < 0.05) for compositionality at some point between 0-1000 ms after the second word; 97 electrodes (green) across 17 patients for lexicality; 3 electrodes (red) for effects of both lexicality and compositionality across 3 patients.

**Figure 4:**
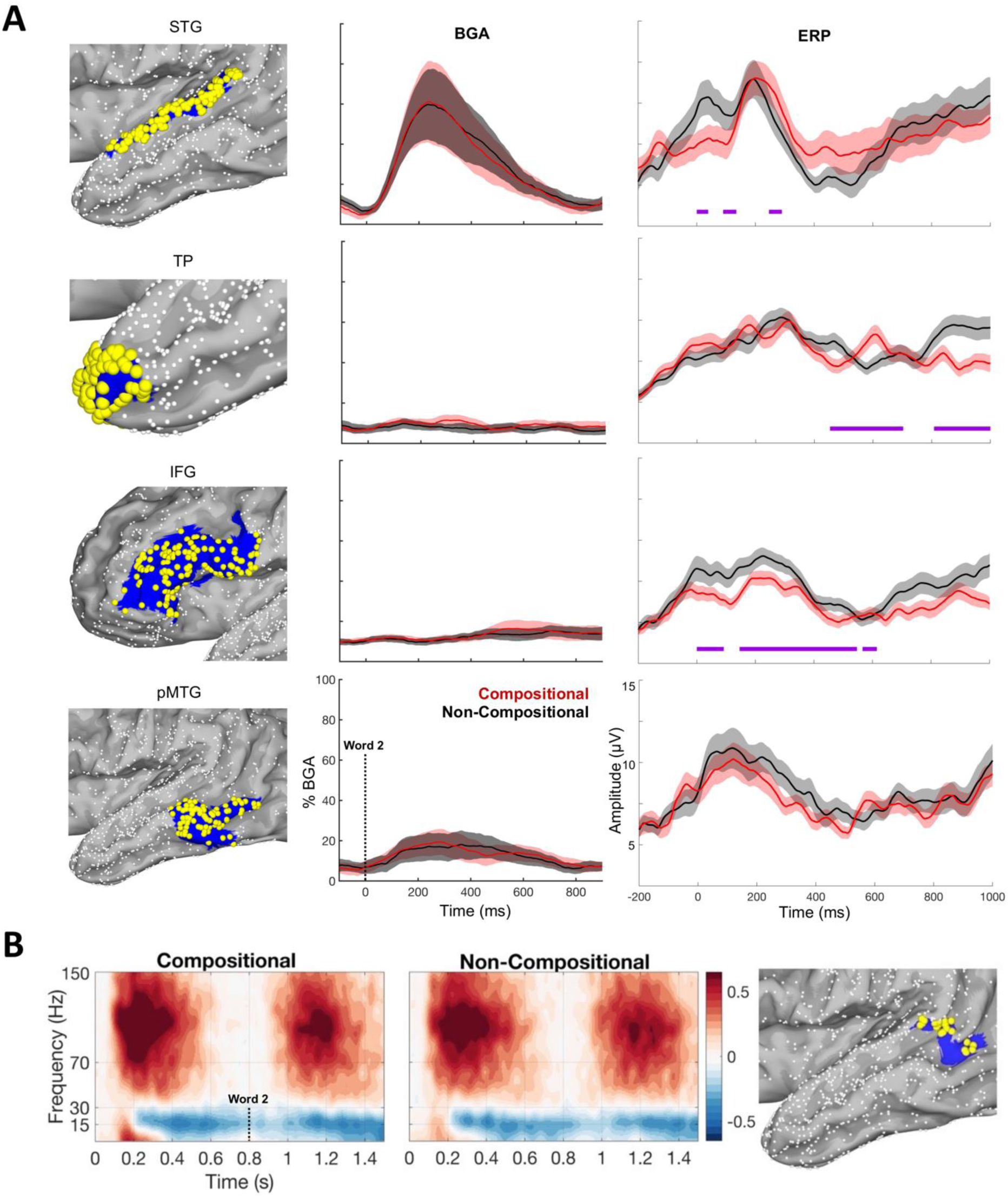
Regions of interest and their broadband high gamma signatures. (A) Regions of interest (left column) with representations of their broadband gamma activity (middle column) and event-related potentials (right column). Red: Compositional. Black: Non-Compositional. HCP index from top-down: posterior superior temporal gyrus (pSTG) = [A4]; temporal pole = [TGd]; inferior frontal gyrus (IFG) = [FOP4, 44, 45, IFSp, p47r, IFSa, 47l]; posterior middle temporal gyrus (pMTG) = [PHT, TE1p]. BGA traces are thresholded by p < 0.05 significantly active from pre-stimulus baseline (−500 to -100 ms) with a minimum of 10% BGA amplitude increase during the 100-400ms window after word 2 onset. ERPs were calculated across the four ROIs plotted on the left column, with significant condition differences being calculated across 0-1000 ms in the same (FDR-corrected) manner as the BGA plots. (B) Spectrograms and electrodes in pSTS for all active channels (HCP index: TPOJ1; electrodes = 23, patients = 10). pSTS electrode coverage per patient: 1.9 ± 1.4 (mean ± SD). Word 2 conset was at 800 ms.

Next, we isolated regions of interest to derive cortical functional connectivity during phrase composition. These were based either on results from our main analysis (pSTS) or on composition effects described in the literature (inferior frontal regions, temporal pole) (Graessner et al., 2021b). To characterize functional connectivity between these regions during phrase composition, we computed phase-locking values (PLV) for electrode pairs situated within pSTS with either pars triangularis or anterior temporal lobe. We computed the generalized phase of the wideband filtered (3–50 Hz) signal that has previously been shown to be more effective than the use of narrowband alpha or theta filters (Davis et al., 2020).

Among patients with concurrent coverage in pSTS and pars triangularis (n = 8, electrode pairs = 231), the majority (n = 5) exhibited significantly greater PLVs for compositional than for non-compositional trials during the 0–500 ms period after second word onset, averaging across PLV values for each pair. In patients with joint coverage in pSTS and temporal pole (n = 8, electrode pairs = 274), the majority (n = 6) showed greater PLVs for the same contrast during the same time window (Fig. 5A, B). We also contrasted PLVs for compositional electrodes and non-compositional electrodes in pSTS, for the subset of patients that had such electrodes. This revealed that compositional electrodes in pSTS exhibited significantly greater PLVs with their paired electrodes in temporal pole and pars triangularis than non-compositional electrodes, with compositional trials also yielding greater peak PLV values. An exemplar patient for pSTS-pars triangularis connectivity is plotted in Fig. 5C.

**Figure 5:**
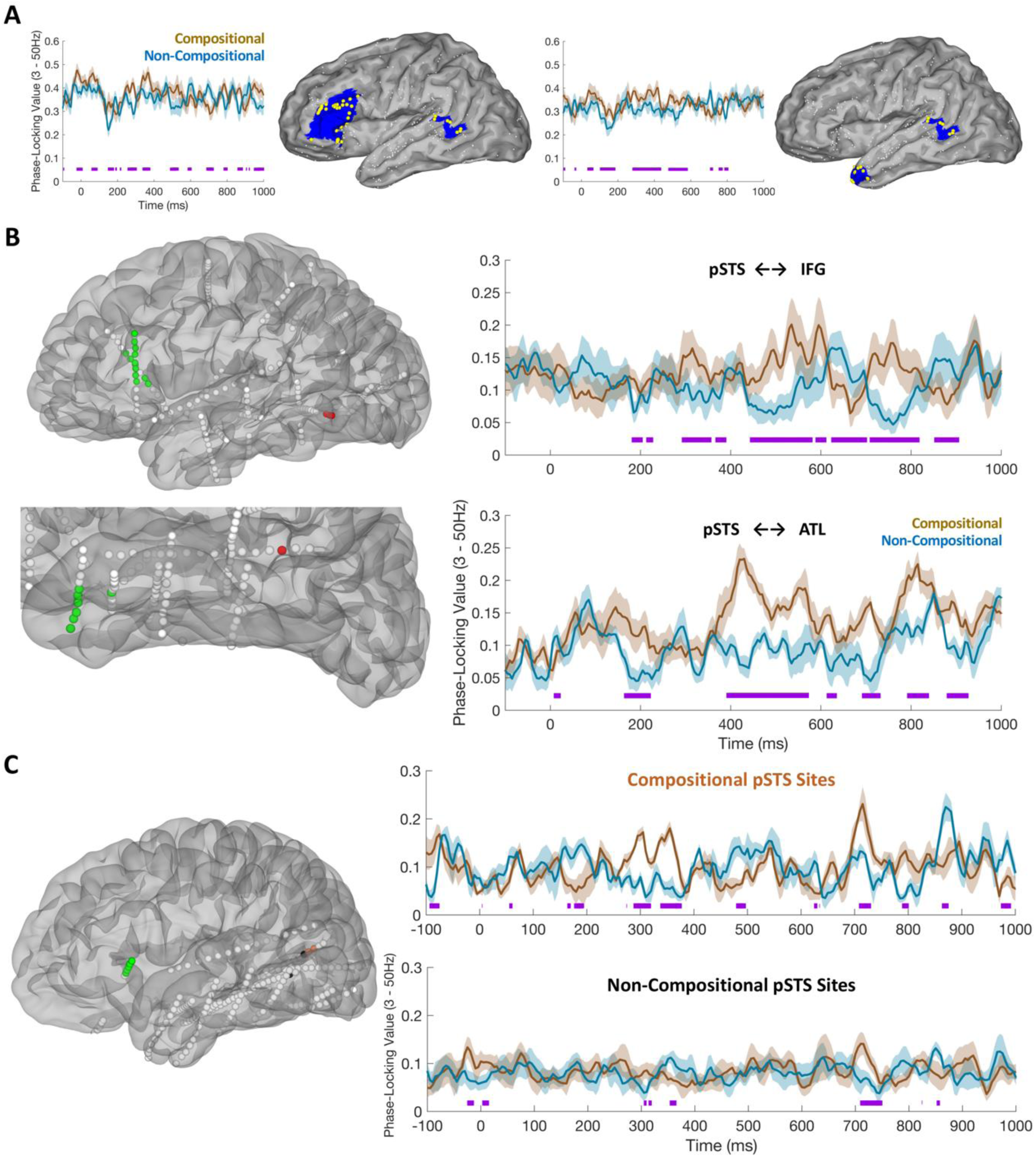
Phase-locking between semantic composition regions of interest. (A) Left: Average generalized phase-locking values (gPLV) for 5 patients showing greater gPLV for phrase composition relative to non-composition between pSTS (HCP index: TPOJ1) and pars triangularis (HCP index: 45, IFSa, IFSp, 47l). Right: Average gPLVs for the 6 patients showing greater phase-locking between pSTS and temporal pole (HCP index: TGd). Purple lines indicate points of significant conditional differences in gPLV values (FDR-corrected for multiple comparisons). gPLV values are plotted from pre-stimulus baseline (−500 to -100ms before first word onset). (B) Posterior temporal lobe (pSTS) gPLVs with inferior frontal gyrus (specifically, pars triangularis) (Top) and anterior temporal lobe (specifically, temporal pole) (Bottom). Left plots show the localization in native space of electrodes significantly involved (q < 0.05) in inter-regional phase-locking (3–50 Hz). Right plots show average time courses (mean ± SEM) of phase-locking value changes from baseline in phrase composition (brown) and non-composition (blue) trials. (C) Phase-locking values between pSTS and pars triangularis in an exemplar patient, contrasting PLVs for electrodes in pSTS that showed an effect of phrase composition (top plot, orange electrodes) with those in pSTS that did not show an effect of composition (bottom plot, black electrodes). Since pSTS is a small ROI, only a subset of our patients satisfied the criteria for this analysis: (1) exhibiting joint coverage across pSTS and pars triangularis; (2) having electrodes in pSTS that did show an effect of composition, and other electrodes that did not.

### Integration of linguistic and visual information

Comparing the Adjective-Noun trials which contained a matching versus non-matching picture, during the 250–500 ms post-picture window SB-MEMAs revealed two notable effects (Fig. 6): The anterior insula (*β* = 0.16; *p* <.001) and pars triangularis (*β* = 0.09; *p* = .002) exhibited greater BGA for phrase-picture matches, while more dorsal frontal regions, centered around inferior frontal sulcus (*β* = 0.10; *p* <.001), exhibited greater BGA for phrase-picture non-matches. Since activity of opercular regions can be misattributed to the insula (Naidich et al., 2004; Woolnough et al., 2019), we ensured that these effects across patients specifically came from electrodes in insula proper by manually checking MRI reconstructions of electrode localizations.

**Figure 6:**
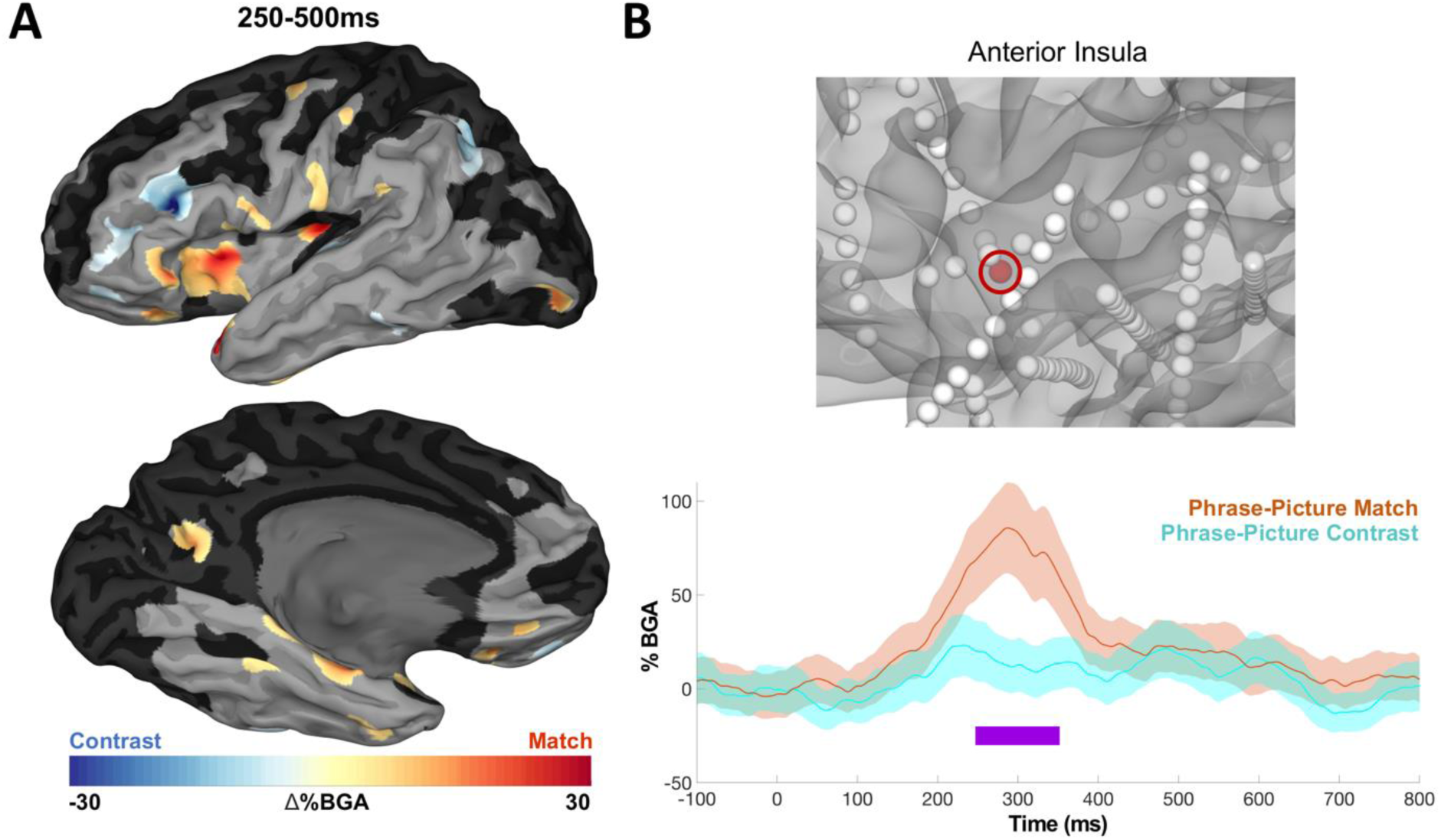
Grouped analysis for linguistic-visual integration. (A) SB-MEMA in BGA for phrase-picture match (orange) and phrase-picture contrast (turqoise) increases, 250– 500 ms after picture onset (threshold: %BGA > 5%, t > 1.96, patient coverage ≥ 3; *p* < 0.01 corrected). (B) Exemplar insula electrode.

## Discussion

We localized the neural signatures of phrase comprehension using minimal adjective-noun phrases in a large patient cohort. We identified a broad portion of posterior temporal cortex as being sensitive to lexicality, and identified neighboring portions in pSTS that respond exclusively to phrase composition. This finely organized heterogeneity in responses implies a cortical topography that takes us beyond traditional structure-function mappings for higher-order syntax-semantics (Naidich et al., 2004). This mosaic architecture has been ascribed to sensory cortices, and to the best of our knowledge has not been shown for higher-level processing in associative cortices (Fox et al., 2020; Tsao, 2020). We speculate that such organization may be a foundational principle underpinning human language. In this section, we discuss these results in the context of previous findings, and evaluate alternative explanations.

Anterior portions of IFG exhibited greater BGA for pseudowords at word 2 position relative to word 1, possibly indexing increased unification demands with the preceding adjective (Hagoort, 2005, 2013). This implies a role for IFG in the unification of more or less expected continuations, in agreement with earlier results (Hagoort, 2004).

Our results address current concerns in the literature with respect to the spatiotemporal dynamics of phrase composition: While IFG activity indexes some aspect of phrase anticipation and phrase-picture matching, it is pSTS that appears to encode the earliest responsiveness to meaningful phrases. These results are supported by lesion-behavior mapping research implicating anterior IFG in executive control for decisions on semantic composition, and broad portions of MTG in representations of individual phrase constituents (Graessner et al., 2021a).

Lastly, anterior insula, pars triangularis and IFS subserve the integration of linguistic input with visual referents. Anterior insula and the IFS have been argued to be the convergence zones of the ventral and dorsal attentional networks (Cazzoli et al., 2021), and our results align with previous models (Willems et al., 2008).

### Lexicality

Our results replicate prior studies in which BGA localized to p/mSTG and pSTS tracks lexicality (Tanji et al., 2005; Humphries et al., 2006; Canolty et al., 2007). This activity may index the bundling of sub-lexical features to yield coherent word-level interpretations of auditory stimuli. Our discovery of greater BGA increases for pseudowords than words at position 2 than position 1 around pars triangularis and surrounding frontal areas (Fig. 1F) suggests involvement of this region in effortful lexical processing to facilitate semantic unification (Hagoort, 2005, 2013). Greater BGA in posterior temporal regions for pseudowords in position 1 may speak to an effect of auditory repetition suppression, or it may indicate greater processing effort as a function of pseudoword phrase position: Greater *lexical access effort* at position 1 (pSTS), and greater *unification effort* (IFG) in position 2.

### Anticipatory response

Based on our results, we suggest that low frequency power in IFG and ATL indexes the preparation of a syntactic slot to be filled by predicted upcoming nominal content. Our findings are in line with the notion that beta oscillations can index the construction and maintenance of sentence-level meaning (Lewis et al., 2016), and the claim that alpha/beta increases can index aspects of syntactic anticipation and category generation (Benítez-Burraco and Murphy, 2019; Murphy, 2020).

### Minimal phrase composition

We found that neighboring portions of pSTS can exclusively code either for lexicality or phrase composition. While a large area of posterior and superior temporal cortex was sensitive to lexicality, a narrower portion was recruited exclusively to code for phrase structure (210–300 ms), generating from simple lexical meanings representations of ‘complex meaning’ (Hagoort, 2020). The same portion of pSTS exhibited greater BGA for non-compositional trials (i.e., pseudowords) at later time points (300-700 ms), implicating it in effortful phrase structure derivations or late-stage lexical search effort/reanalysis. In addition, the greater BGA for phrase composition in pSTS was crucially found across different electrodes to those sites that showed greater BGA for late-stage pseudoword processing, which we believe contributes to a cortical mosaic. ERP responses were also dissociable from BGA: A late effect for non-composition was found in Broca’s area, likely due to greater attempted lexical access for pseudowords, while a late signature was detected in temporal pole for composition, potentially related to late-stage conceptual access.

Phrase composition was also marked via increased functional connectivity between pSTS electrodes implicated in composition and both inferior frontal regions and temporal pole. This appears to reflect network-level interactions seemingly required for basic phrase formation (Baggio and Hagoort, 2011; Schoffelen et al., 2017). While the duration of composition effects lasted approximately 500 ms, there were nevertheless also late-stage periods of brief (∼80 ms), reversed effects. As such, future intracranial research involving either more dense or fine-grained coverage could provide a clearer picture.

Anatomical connectivity has been elaborated by white matter pathways that connect pSTS with IFG and ATL (Glasser and Rilling, 2008; Figley et al., 2017; Sarubbo et al., 2020). We theorize that the formation of phrase structure in pSTS feeds categorial information to conceptual interfaces in ATL and memory/control interfaces in pars triangularis. One such piece of information would be the phrase category/label.

Previous intracranial research found that the lower bank of the STS is involved in lexico-semantic processing (Nourski et al., 2021), and our results appear to suggest greater involvement of the lower bank of the pSTS in semantic composition than the cytoarchitechtonically distinct upper bank (Zheng et al., 2010). Future work utilizing even higher scale recording techniques, such as single-unit recordings (Bitterman et al., 2008), will more comprehensively address this issue.

One of the main candidates for phrase composition from previous research, ATL, was implicated via late-stage ERPs. ATL activity has been found to be delayed until after intra-lexical morphological composition (Flick et al., 2018). This implies the existence of composition-related activity independent of semantic and orthographic processing. Our findings suggest that pSTS might be one such region, although since we only presented auditory stimuli we cannot make any stronger claims.

Our findings are concordant with recent results and models implicating the pSTS in phrase composition (Nelson et al., 2017; Flick and Pylkkänen, 2020; Matchin and Hickok, 2020; Murphy, 2020; Law and Pylkkänen, 2021; Matar et al., 2021); producing hierarchically organized functional morphemes (Lee et al., 2018); sign language comprehension (Trettenbrein et al., 2021); cross-linguistic reading competence (Feng et al., 2020).

Existing models have posited distinct neural sites across gross portions of the cortical mantle for lexicality and basic composition. Typically, this is a large scale “posterior vs anterior temporal” (Friederici, 2012) or “dorsal vs ventral stream” distinction (Bornkessel-Schlesewsky and Schlesewsky, 2013), or a distinction between ATL (conceptual-semantic) and pMTG (hierarchical structure) (Matchin & Hickok, 2020). In contrast, we report overlapping functionality at a much smaller scale in the pSTS.

We have claimed that pSTS is “tiled” in a mosaic-like structure coding for lexicality and phrasal meaning. However, there is an alternative account invoking pseudoword and task-related processing effort. We believe that it is likely that our late-stage (300-700 ms) signature of greater BGA for non-compositional trials in pSTS can be explained by general difficulties with pseudowords. Yet, the effort-related account seems unrelated to our early (100-300 ms) BGA increases for compositional trials; we found no clear posterior temporal differences for real words in non-compositional trials, which the effort-related account would predict (due to nouns being presented after pseudowords); patients that showed early BGA increases for composition also showed distinct signatures of non-composition processing effort in neighboring (but distinct) sites in pSTS and surrounding cortex (Fig. 3). A comparison of real word processing in the non-compositional trials (Fig. 1G) revealed no clear effects in posterior temporal regions, only a BGA increase for nouns over frontal opercular sites, suggesting any degree of task-related effort was isolated to late-stage pseudoword processing.

We have provided a more spatiotemporally reliable profile of basic semantic composition than previous non-invasive methodologies, resolving some conflicts in the literature concerning the variety of effect timings documented. In addition, we believe that our results point to the need to move beyond simpler hypotheses about pSTS either being involved in phrase composition or not: Instead, it appears to be a question of *when* and *over which specific portions of tissue* (e.g., on the order of 10mm). We believe that recordings with higher spatial resolution may reveal greater insights into the operations of this cortical mosaic that we have documented.

While our paradigm only allows us to make direct claims about semantic composition, we suspect that portions of pSTS are also implicated more narrowly in minimal syntax, however the functional connectome of the syntax network may differ from that of the semantic compositional network. Given that recent research has revealed substantial overlap in the distribution of the fronto-temporal language network for speakers of 45 languages across 11 language families (Ayyash et al., 2021), and given that the computational process our paradigm isolated was generic, we also expect that pSTS activity codes for meaningful phrase structures across languages other than English. The specific neural signatures of basic phrase structure composition isolated here represent an elementary computation in comprehending natural language that can be probed further in future intracranial research.

## Acknowledgements

We express our gratitude to all the patients who participated in this study; the neurologists at the Texas Comprehensive Epilepsy Program who participated in the care of these patients; and the nurses and technicians in the Epilepsy Monitoring Unit at Memorial Hermann Hospital who helped make this research possible. This work was supported by the National Institute of Neurological Disorders and Stroke NS098981. Human subjects: Patients participated in the experiments after written informed consent was obtained. All experimental procedures were reviewed and approved by the Committee for the Protection of Human Subjects (CPHS) of the University of Texas Health Science Center at Houston as Protocol Number: HSC-MS-06-0385.

